# Systematic Investigation of Double Emulsion Dewetting Dynamics for the Droplet Microfluidic Production of Giant Unilamellar Vesicles (GUVs) under Biocompatible Conditions

**DOI:** 10.1101/2025.02.04.636495

**Authors:** Wenyang Jing, Heewon Noh, Timothy J. C. Tan, Nicholas C. Wu, Hee-Sun Han

**Affiliations:** Center for Biophysics and Quantitative Biology, University of Illinois Urbana-Champaign, 600 S Mathews Ave, Urbana, Illinois 61801, USA; Department of Chemistry, University of Illinois Urbana-Champaign, 505 South Mathews Ave., Urbana, Illinois 61801, USA; Department of Chemical and Biomolecular Engineering, University of Illinois Urbana-Champaign, 600 S Mathews Ave., Urbana, Illinois 61801, USA; Department of Biochemistry, University of Illinois Urbana-Champaign, Urbana, IL, 61801, USA; Carl R. Woese Institute for Genomic Biology, University of Illinois Urbana-Champaign, 1206 W Gregory Dr, Urbana, IL 61801, USA; Carle Illinois College of Medicine, University of Illinois Urbana-Champaign, 807 South Wright St., Urbana, Illinois 61801, USA

## Abstract

Giant unilamellar vesicles (GUVs) embody biomimetic membranes with compartmentalization that serve not only as simplified models to better understand complex biochemical and biophysical processes, but also as a chassis for the bottom-up assembly of synthetic cells. Recently, double emulsion droplet microfluidics has proven to be a promising platform for their production, offering greater throughput, control, and reproducibility over traditional methods. However, the interplay of parameters—particularly under biocompatible conditions—that influence the complex multiphase fluid dynamics of the dewetting process underlying GUV production has not been thoroughly studied, limiting the democratization of the approach. In this study, we systematically investigate how lipid composition and concentration, aqueous phase conditions, droplet confinement, and fluid dynamics effects promote or impede the dewetting process. We show that the prevalent use of high concentrations of glycerol and P188 are unnecessary, and the altered dewetting dynamics with restricted surfactant usage can be tuned by adjusting chip dimensions and multi-phase compositions. Guided by these findings, we achieved robust, high throughput production of monodisperse GUVs using 0.1% P188 and no glycerol with salts. Our results improve the reliability and accessibility of droplet-microfluidics GUV platforms to catalyze advances in biophysics, synthetic biology, and drug discovery.

## Introduction

The fundamental unit of life, the cell, requires multiple signaling pathways, interacting components, and regulatory networks to operate, thus embedding inherent complexity that can complicate efforts to understand basic functions or to evaluate drug efficacy. In order to simplify and attempt to isolate systems of interest for study, the traditional top-down or reductionist approach has been employed to try and create the minimal cell, but genes and proteins of unknown function remain^1,2^, therefore leaving potentially undesirable influence and interference in the system. Conversely, the bottom-up approach^3–5^ promises a controlled way to include only the known and desired components of interest. This has led to the idea of the artificial or synthetic cell^5–10^, which not only provides a model to facilitate biochemical and biophysical studies^11–15^ but also offers the potential for bioengineering and biomedical applications^5,16–20^. To this end, giant unilamellar vesicles (GUVs)^9,11,15,21,22^ are the most attractive chassis for building the synthetic cell owing to its lipid bilayer and enclosed compartmentalization that together serve as a biomimetic host for natural or engineered functions. Traditionally, GUVs have been made using bulk approaches^11^, but they have poor throughput, encapsulation efficiency, and reproducibility. In contrast, droplet microfluidics overcomes these deficiencies and is perhaps the best approach for templating GUVs^23–26^, offering improved control over size, cargo encapsulation efficiency, production throughput, and fluid environment.

In particular, double emulsion (DE)-based approach, which utilizes an inner aqueous (IA) phase, a middle lipid-oil (LO) phase, and an outer aqueous (OA) phase, has gained increasing traction^16,26–33^. This approach leverages the inherent solubility of lipids in oils to create a barrier between the two aqueous phases, mimicking cytosolic compartments (IA) and extracellular fluids (OA), respectively. The LO phase spontaneously dewet or phase separate, leaving behind a lipid bilayer. Compared to the single emulsion (water-in-oil)-based approach^25^, which relies on surfactant-stabilized droplet interface for liposome formation and GUV release through droplet break-up, the DE-templated method is gentler and provides superior throughput and monodispersity in GUV size.

Previously, oleic acid^34^ or a mixture of chloroform and hydrocarbons^35–37^ were utilized as the oil phase of DE, but it was recently demonstrated that octanol or octanol mixtures show faster kinetics and enhanced biocompatibility^33^. However, the increasingly adopted octanol approach includes Poloxamer 188 (P188)—also known as Pluronic F-68— and glycerol to efficiently drive phase separation, or dewetting, and to protect membrane integrity at high surfactant concentration, respectively. P188 is included at 5-15%^16,26–31,33^ and glycerol concentration is kept at 15%^16,26–29,33^, which increases membrane viscosity by several orders of magnitude^38^. Unfortunately, these conditions are incompatible with enzymes or membrane proteins^39–41^, compromising the potential of GUVs as functional synthetic cells. For example, such high concentration of glycerol increases membrane viscosity^38^, affects membrane protein functions^39–41^, permeates the membrane and disturbs internal biochemical reactions^31^, and decreases gas permeability of the membrane^42^. Indeed, the commercially available *E. coli* S30 in vitro transcription/translation kit notes that glycerol >1% is inhibitory^43^. Although P188 is a known cell media additive in bioreactors either as a shear protectant or to reduce cell adhesion^44^, 5% concentration represents 50 times the typical level used in the cell culture. At this high concentration, P188 has multiple negative impacts on protein function, including inhibition of protein polymerization^26^ and induction of protein aggregation even at lower concentrations^45^. In this study, we show that these limitations extend to membrane proteins. Using SARS-CoV-2 spike protein and Angiotensin-converting enzyme 2 (ACE2) protein stably expressed on mammalian cells as models, we show that membrane protein function is compromised even at 1% P188 (Figure 2). Due to these limitations, GUVs produced via the DE-based method have been predominantly used in applications involving a limited set of proteins, such as alpha-hemolysin^26^ and soluble proteins^26,46^, rather than complex, functional membrane proteins.

To further enhance the versatility of DE-based GUV production, we aim to establish biocompatible conditions, especially with minimal P188 and no glycerol. While these parameters address biocompatibility concerns, other questions of dewetting dynamics remain. Recent studies have found that changes to the aqueous conditions can result in different lipid concentration requirements and flow conditions for dewetting^30^ as well as the inhibition of dewetting by salt^46^. To date, it is still unclear not only how best to choose experimental parameters to achieve the desired dewetting outcomes but also how to achieve dewetting in fully biocompatible conditions.

To address these uncertainties, this study presents the first systematic investigation into how different parameters impact dynamic dewetting behavior, providing insights into DE-templating GUV production. We developed a two-stage microfluidic chip and/or centrifugation that allows precise control over experimental conditions. We independently varied lipid compositions, including charge, molar ratios, and acyl chain saturation as well as P188 and lipid concentrations. The fluid dynamic effects such as on-chip flow speed and centrifugal force, and confinement were also varied. We establish how these parameters are correlated with dewetting success and show how our study design and findings offer a practical means to increase throughput and reliability. Overall, we show that the dewetting process is complex, driven by fluid dynamics under confinement, with surfactants dynamically redistributing around the interface. This behavior cannot be fully explained by thermodynamic perspectives of traditional static models. For biocompatibility, we demonstrate that glycerol is unneeded, that the well-established biocompatibility of 0.1% for P188 (50x lower) is feasible, and that it is also possible to induce dewetting of stable GUVs with biorelevant lipid, surfactant, and salt/buffer conditions. These results offer insights on the physical process and provide design guidance to facilitate the adoption of the droplet-microfluidic GUV approach. By improving its accessibility and usability, this approach can better harness the advantages of GUVs, unlocking their full potential for basic research and a wide range of life science applications.

## Results and Discussion

### Theory on DE dewetting and tested parameters

On-chip DE-templated GUV production involves two steps: DE droplet generation and the subsequent phase separation of DE droplets under flow, in contact with confined walls (Figure S1). During phase separation, DE droplets transition into an intermediate state, where the oil pocket shifts to the side and the lipid bilayer zips off between aqueous phases. If the oil pocket continues to protrude outwards and eventually splits off, it forms GUV. In our study, three outcomes were observed depending on the degree of dewetting: 1) not dewetted (remaining as DE), 2) partially dewetted (PD), 3) fully dewetted (GUV) as shown in Figure 1A.

**Figure 1.**
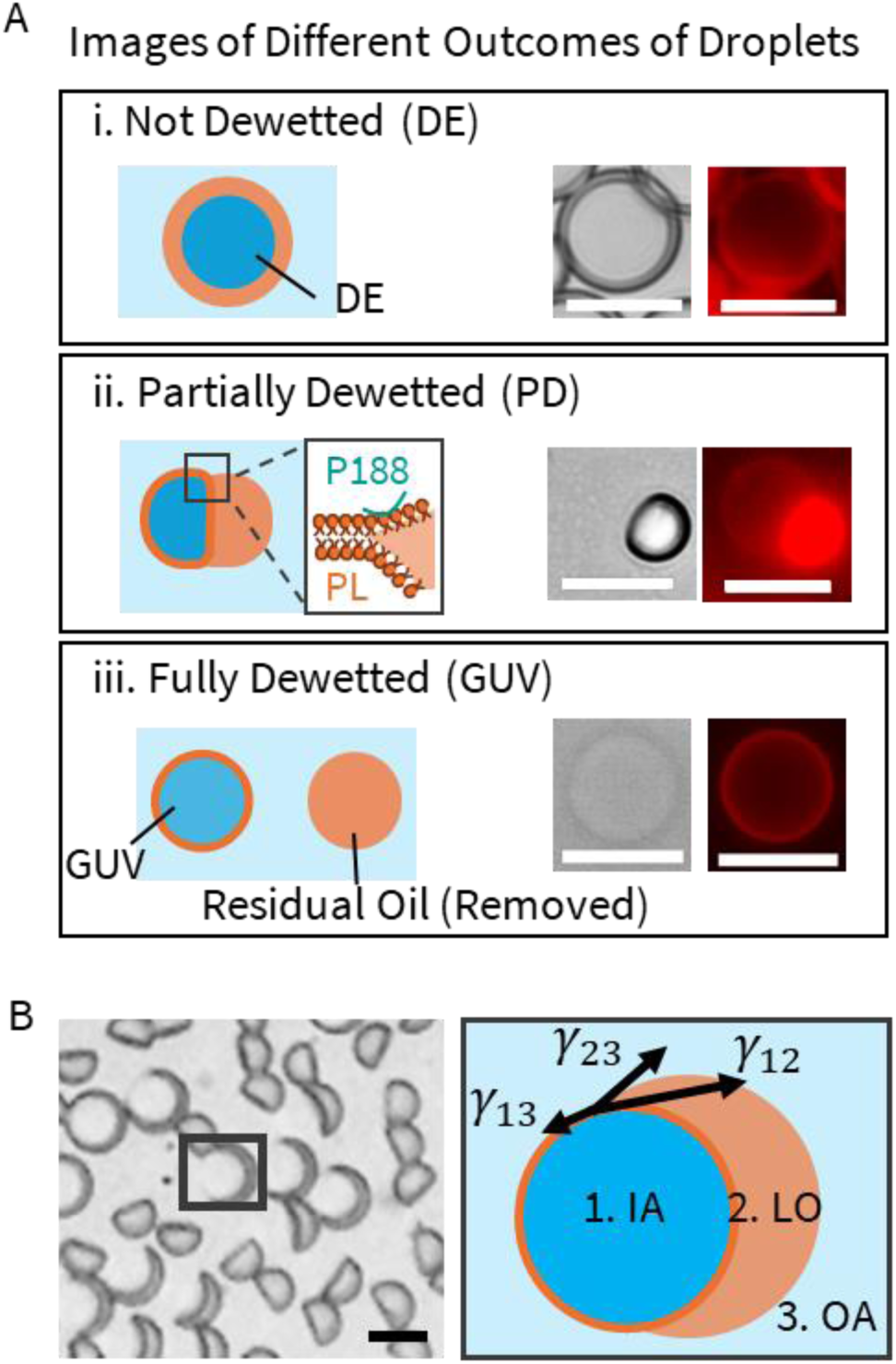
Interfacial tension affects dewetting. (A) Three outcomes of dewetting: (i) double emulsions with oil shells, (ii) partially dewetted vesicles with side-attached oil pockets, and (iii) fully dewetted GUVs, characterized by poor contrast in brightfield image. Each panel includes an illustrative demonstration on the left and brightfield and fluorescence images on the right. GUVs are labeled with 0.2% Liss Rhod PE, a fluorescent lipid dye. P188 refers to Pluronic F68, and PL denotes phospholipids. Scale bar: 20 µm. (B) Three interfacial tension values involved in the dewetting of DE droplets. DE droplet in the highlighted frame is shown as an example. Scale bar: 20 µm.

**Figure 2.**
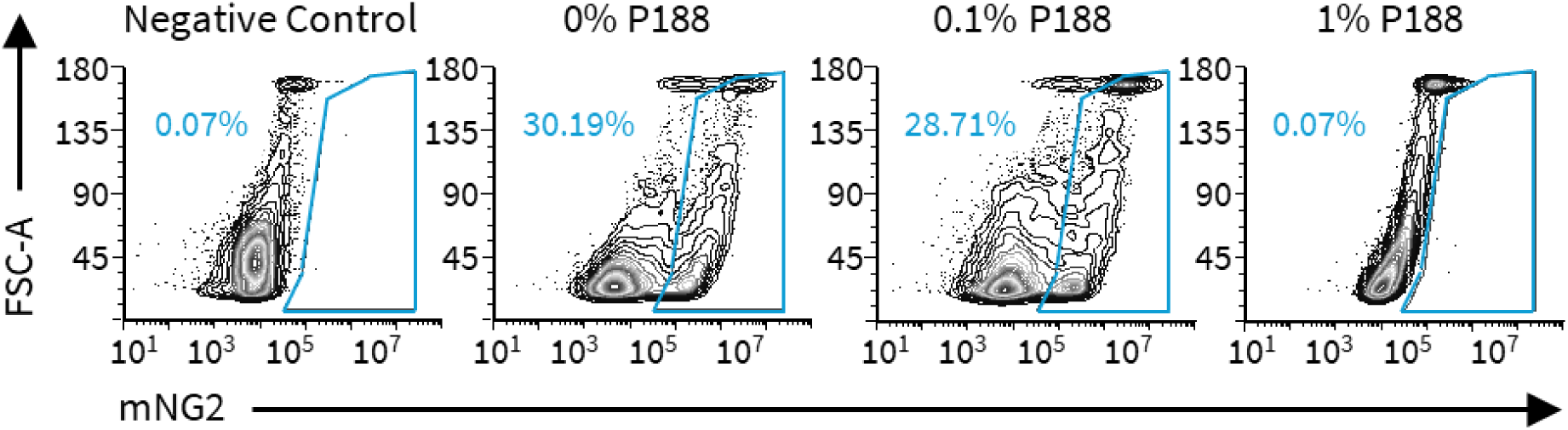
Flow cytometry-based detection of fluorescence of syncytia based on recombined mNG2, whose intact form is produced when the cytosols of two different engineered cell lines successfully fuse, with each type expressing one part of mNG2 intracellularly, and either ACE2 or SARS-CoV-2 spike on the cell surface (see Materials and Methods). Percentages indicate the fraction of cells that have fused. Note that the negative control, which was a co-culture utilizing ACE2 cells that did not express one part of mNG2, and therefore did not generate fluorescence upon cell-cell fusion, shows the same lack of detection as the 1% P188 case. In comparison, there is only a slight decrease from ∼30% of the positive control (0% P188) to ∼29% for 0.1% P188.

**Figure 3.**
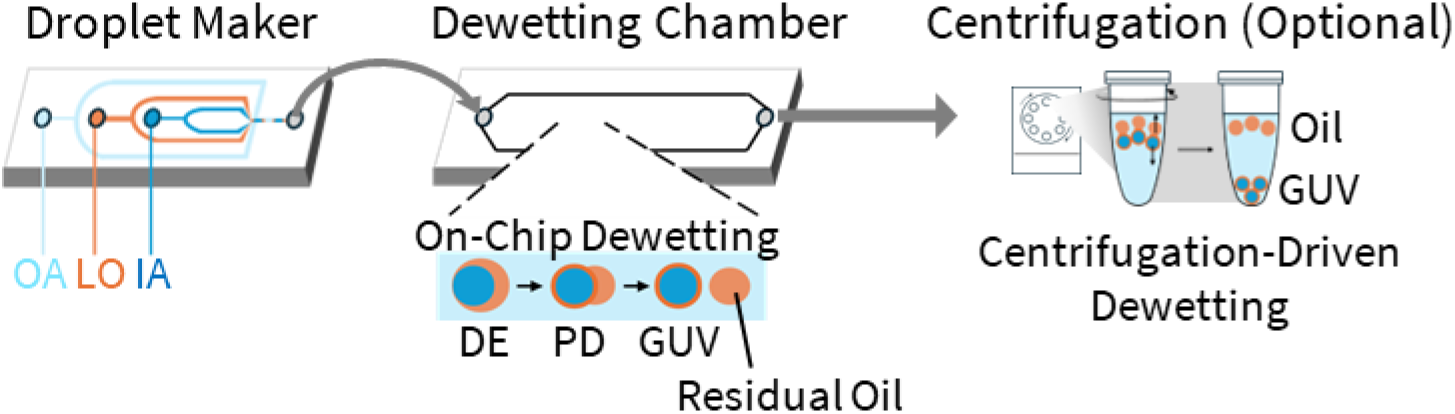
Illustration of microfluidic and centrifugation setup used to investigate dewetting dynamics. A droplet generator and dewetting chip are used to decouple droplet making from downstream flow conditions during dewetting and to decouple surface treatments. Centrifugation is used in cases of partial dewetting on-chip, enabling partially dewetted droplets to fully dewet by leveraging the density difference between aqueous and oil layers. DE and PD refer to double emulsion droplets and partially dewetted vesicles, respectively.

Conventionally, the dewetting process for DE droplets is explained by surface energy minimization, described by the spreading coefficient^37,50,51^, which depends on the three surface tensions involved. Figure 1B illustrates how in the partially dewetted state, these three interfacial tensions interact at the contact point. The spreading coefficient is defined as:

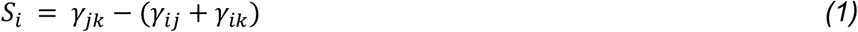

where phases are defined as follows: IA (inner aqueous phase): 1, LO (lipid oil phase): 2, and OA (outer aqueous phase): 3. For dewetting to occur, the following three inequalities must be satisfied.

1. The LO phase must resist spreading over the other phases

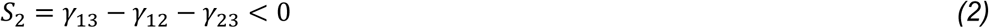
2. The OA phase must favor spreading over the other phases, not IA

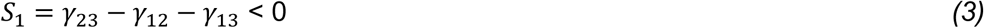

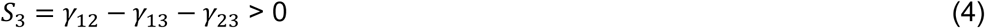

These conditions drive phase separation of the oil phase, causing the oil droplets to bud, leaving a lipid membrane between the aqueous phases. Complete dewetting occurs when the surface tension between the IA and oil (γ_12_) exceeds both the interfacial tension between the OA and oil (γ_23_) and the lipid bilayer tension (γ_13_). Since the lipid bilayer tension is in the order of µN m^-1^, two to three orders of magnitude lower than the surface tension between the oil and aqueous phases ^52,53^, the key requirement is ensuring the surface tension γ_23_ remains smaller than γ_12_. This has been achieved using a higher concentration of P188 in the OA phase. However, significantly reducing the P188 concentration to a biocompatible level will substantially increase γ_23_ thereby hindering phase separation.

All three interfacial tensions are determined by not only concentrations of P188 in the aqueous phases but also the types and concentration of lipids in the LO and the presence of salts, all of which were systematically investigated in this study. For lipids, we initially use the model lipids DOPC (neutral) and DOPG (negative) in 2:1, 1:1, and 1:2 molar ratios while varying their concentrations from 5 to 35 mg/mL. We measured the surface tension between the LO and aqueous phases (γ_12_ and γ_23_) using pendant droplets (Table S3 and S4). As expected, we found that increasing surfactant and lipid concentration decreases the surface tensions between the LO and both aqueous phases. Additionally, we observed that an increase in lipid negative charge further decreases these surface tensions while adding salt generally increases them. As we could not measure the surface tension between the two miscible IA and OA phases, and some aqueous-LO tensions were too low to form a proper pendant drop, we could not calculate the spreading coefficient. The value of γ_13_ is expected to be low given known values of interfacial tensions between aqueous two-phase systems, which have a typical range between 0.001-0.1 mN/m^52,53^.

We note that although global surface energy minimization predicts the equilibrium states after the system reaches thermodynamic equilibrium, it does not describe the evolution process from non-equilibrium. The existing models assume a stationary DE and not one undergoing time-dependent shear forces and deformation, as is in the on-chip dewetting device.

Additionally, dynamic viscosity changes^54^ and fluid flow^55–57^ could significantly affect the process, indicating that these dynamic parameters can even affect the final state. Thus, we also consider confinement and fluid flow as factors that can alter the timescale and expected state of dewetting.

Our investigations indeed show the strong influence of confinement and flow rate on the dynamic dewetting process, underscoring the importance of exploring dynamic factors. These results are consistent with the previous simulation study on vesicles showing that high confinement drives lipid domain reorganization thus altering the membrane tension distribution. This observation suggests that the interfacial molecules of both lipids and P188 may undergo dynamic distribution changes under flow and confinement. The redistribution of surfactants creates a dynamic surface tension gradient, generating tangential stress along the interface (Marangoni)^59^, that can influence the separation force and compromise the lipid membrane integrity. To evaluate the strength of Marangoni effects, we use the Marangoni number, *Ma* = Δγa/(ηD), where γ is the interfacial tension, a is the droplet diameter, η is the dynamic viscosity, and D is the diffusion coefficient. The diffusion coefficient of P188^60^ and monolayer lipids^61^ are similar and of order 10 µm^2^/s, whereas lipids in a bilayer have a diffusion coefficient that is an order of magnitude lower^33^. For a 10 µm droplet with a mean surface tension of 0.01 mN/m, even a 1% difference for Δγ yields an estimated Marangoni number of ∼100, which indicates the possibility of strong Marangoni stresses. This analysis demonstrates how various parameters can intricately influence dewetting through Marangoni-driven flows as well as interfacial shear.

In summary, a thermodynamic perspective is insufficient to predict dewetting outcomes within practical timeframes. As will be shown in the following section, certain conditions resulted in partial dewetting, where external forces, such as centrifugation, helped to achieve complete dewetting, suggesting a metastable state rather than thermodynamic equilibrium. Additionally, multiple parameters interactively affect the dewetting status, exhibiting reversed trends depending on other conditions. This complex trend underscores the need for a systematic investigation of each factor. In this study, we systematically examine how different experimental parameters impact the dewetting process for GUV production. Table 1 summarizes the parameters that are tested.

**Table 1.** Tested Parameters were summarized in table. IA, LO, OA represent the inner aqueous, lipid in oil, and outer aqueous phases.

| Dewetting Parameters Tested in This Study |
| --- |
| P188 Concentration in OA |
| Lipid Charge in LO: DOPC(neutral)+DOPG(negative) |
| Lipid Concentration in LO |
| Salt and buffer in IA and OA |
| Confinement: DE Diameter/Chamber Height |
| Flow Velocity on Dewetting Chamber |
| Lipid Acyl Chain Saturation in LO (last section) |

### Bioincompatibility of previously used conditions for DE-based GUV production

For glycerol and P188, our choices stem from biocompatibility and dynamics considerations. We aim to exclude glycerol from the system as it can bind to the membrane altering their properties and pass through the membrane therefore interfere with IVTT reactions^43^. Dialysis-based removal of glycerol has been implemented, but it is labor-intensive^62^. The absence of glycerol will offer new insights into dewetting dynamics while its influence as a viscosity modifier can still be contextualized and deduced. For P188, we aim to use 0.1% P188, a concentration equivalent to 1x concentration used in cell culture, which is 50 times lower than what has been used to promote dewetting.

Building on the knowledge that P188 prevents cell adhesion to air bubbles through insertion or adsorption into the cell membrane^44^, our study reveals that even at 1% P188, 5 times lower than the concentration commonly used for GUV dewetting, some cell functions and associations are impaired, emphasizing the need for a significant reduction in P188 concentration. Specifically, we tested the cell-cell fusion activity mediated by interactions between membrane-bound ACE2 and SARS-CoV-2 spike protein (Figure 2). In this assay, we used two engineered HEK293T cell lines each stably expressing a complementary fragment of split fluorescent protein mNG2^63^ intracellularly, and either membrane-bound ACE2 or SARS-CoV-2 spike on the cell surface, mimicking viral entry into cells^49^ (see Methods). The effects of P188 on ligand-binding mediated cell-cell fusion are significant as shown in Figure 2. At 0% P188, the successful cell to cell fusion rate is 30.19%, while 1% P188 completely inhibits cell-cell fusion (0.07%). At the commonly used 0.1% concentration, the fusion rate is only slightly lower than that in the absence of P188. This result aligns with the previous report suggesting that P188 may directly bind to SARS-CoV-2 spike protein, blocking viral entry into cells, as demonstrated in viral infection assays and molecular dynamics (MD) simulations^64^. Additionally, P188 can be incorporated into cell membranes and insert into membrane pores as a resealing agent, which may prevent fusion^65^.

Even with the subsequent washing step, using high P188 is also undesirable as complete removal is a noted difficulty^44^, with P188 being known to adsorb and insert into the lipid bilayer^44,66,67^. As an example, the potential for P188 effects to persist has been documented in terms of preventing cell adhesion, where it took 6 to 7 passages in 12 mL cell flasks to recover a stable level of cell adherence from a starting concentration of 0.1% P188^44^. If one generously assumes 10% of previous cell media is left behind each passage, it would mean that a minimum dilution of 10^6^ to 10^7^ would be required. A comparison can also be drawn with the surfactant Triton-X 100, which has a critical micelle concentration (CMC) of 0.22 mM, similar to that of P188 (0.48 mM); it is known that even prolonged dialysis is insufficient to remove low CMC surfactants like Triton-X from the lipids they associate with during proteoliposome formation^68^.

### Microfluidic Chip Design

We designed a two-stage system comprising a separate droplet maker and dewetting chamber connected by PE tubing. In this setup, DE droplets are generated in the first chip and undergo phase separation in the second. This two-stage configuration offers several advantages over conventional single-stage chips^26^. First, it enables straightforward, single-layer fabrication of each component with distinct heights, eliminating the need for aligning different PDMS layers to create a multi-height device. Second, it allows for separate surface treatments of the droplet maker and dewetting chamber, streamlining fabrication, and significantly increasing success rate. Producing double emulsions requires a surface wetting change at the droplet-making junction from hydrophobic to hydrophilic. In the one-stage system, controlling polymer deposition at this junction is challenging because the channel after the junction is wide and long to facilitate dewetting, and polymer is typically introduced from the outlet. A separate droplet maker enables the use of simple plasma treatment-based polymer deposition for the droplet making channel, improving control and precision.

The design of our two-stage chip, combined with biocompatible conditions, improves GUV production throughput by orders of magnitude. Since the dimensional and speed requirements for high-throughput droplet generation and efficient dewetting differ, separate designs for each process facilitate throughput optimization by appropriately scaling dewetting chamber geometry. Moreover, eliminating glycerol and reducing the concentration of P188 further enhance throughput by decreasing the capillary number (*Ca* = ηU/γ, where η is the dynamic viscosity, U is flow speed, and γ is the interfacial tension). This reduction can suppress jetting at the droplet-making junction, which is a critical factor for achieving small channel dimensions required to produce 10–20 µm GUVs.

In the original study^26^, the IA flow rate is estimated to be at most at 1 µL/h, consistent with the estimation from another study^30^, orders of magnitude slower than common droplet microfluidics operation. Prolonged operation at such low throughput is undesirable as octanol interaction with the walls decreases hydrophilicity and their buildup on the surface often clogs the channels^69^. Furthermore, such low flow rates are not practical for typical syringe pump systems. In this study, we operated droplet maker at significantly higher flow rates (IA/LO/OA: 15/15/40 µL/hr), though this is not upper limit. The maximum on-chip residence time was ∼30 s, an order of magnitude lower than previously reported^26^ and sufficient for examining parameter correlations. For future applications, throughput and residence time can be optimized.

For systematic investigation of flow speed and confinement effect on the dynamic dewetting process, we modulated the flow velocity and the channel height by altering the dimensions of the dewetting chamber with fixed flow rates. Compared to the original study^26^ (∼0.5 mm/s), three average flow velocities were tested and spanned an order of magnitude from ∼1 mm/s to ∼20 mm/s, with the corresponding Reynolds numbers (*Re*) ranging from ∼0.03 to ∼ 0.5, where *Re* = ρU*L*/η, ρ is the fluid density, U is flow speed, *L* is the hydraulic diameter of the channel, and η is the dynamic viscosity. Additionally, we examined three different confinements—0.5, 0.8, and 1.2—defined as a/*H*, where a is the droplet diameter and *H* is the height of the dewetting channel.

Centrifugation, an external high-speed flow system, is applied to some conditions to further assist the dewetting^31^ as octanol (0.82 g/mL) is less dense than the aqueous phase (1.04 g/mL). To assess the centrifugation-driven dewetting dynamics, partially dewetted droplets collected from the dewetting chamber were subjected to benchtop centrifugation at three different forces: 2000 × *g* for 1 min, 10000 × *g* for 1 min, and 17000 × *g* for 1 min. By contrast, direct centrifugation of DE droplets resulted in prolonged dewetting, which is undesirable due to the increased risk of GUV fragmentation during extended centrifugation. While increasing the centrifugal force or the duration should enhance the separation, our observations revealed that excessively high mechanical stresses caused GUV fragmentation (Figure S10), consistent with prior studies^70^. We carefully optimized the centrifugation conditions to balance effective dewetting with minimal GUV fragmentation.

### Dewetting Dynamics at High OA Surfactant Concentration Without Glycerol

Previous studies^16,26,31,33^ demonstrated successful on-chip dewetting and GUV formation at 5% P188 and glycerol. We were also able to replicate this result (Figure S7). Eliminating glycerol at high P188 concentrations drove complete dewetting but failed to form intact GUVs. DE droplets transitioned into free-standing octanol droplets accompanied by high fluorescent background signals indicate vesicle rupture (Figure S8). Glycerol likely dampens tension gradients and the overall stress/strain response of GUVs by significantly increasing solvent and membrane viscosities significantly^38^, enabling the intact dewetting of GUVs at 5% P188.

We performed systematic investigation of the dewetting dynamics in 5% P188 without glycerol in OA. This composition, however, failed to form stable DE droplets at the droplet making junction. To address this, we added an inlet downstream of the droplet-maker to introduce and homogenize additional P188 with the DEs before entering the dewetting chamber (Figure S6). Additionally, we systematically test both salt-free and salt-containing (10 mM HEPES with 100 mM NaCl) conditions to evaluate the effect of salt on dewetting dynamics.

At high P188 and no glycerol conditions, the outcomes were categorized into two groups: 1) dewetted & destroyed on-chip: all DEs dewetted and destroyed within the chamber or 2) dewetted & destroyed off-chip: remaining DE droplets spontaneously dewetted within an hour after exiting the chamber (Figure 4). In all conditions, the height of the dewetting channel must be smaller than the outer diameter of DE droplets (confinement: > 1) to achieve on chip dewetting. This result suggests that droplet-wall contact enhances internal circulation through droplet tank-treading, consistent with previous study showing that internal circulation speeds up the transition of the completely engulfed or core-shell state to the partially dewetted state^71^. The following sections explore how other parameters affect dewetting dynamics at the highest confinement, unless specified otherwise.

**Figure 4.**
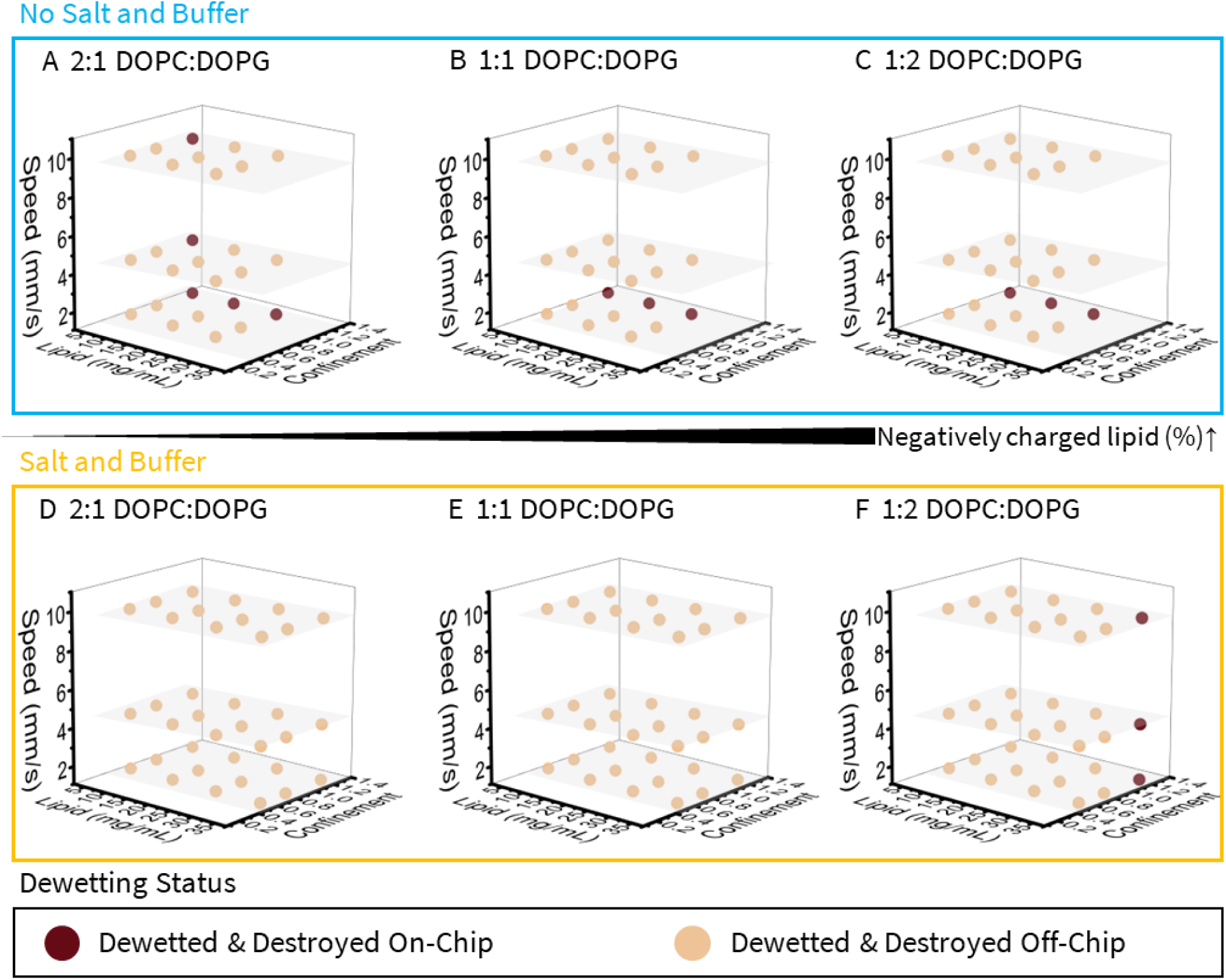
Dewetting results at high OA surfactant concentration (5% P188). These plots show the results of double emulsion dewetting for different negative charge ratios, lipid concentrations, on-chip flow speeds, confinement, and the presence or absence of salt and buffer in the OA (see Materials and Methods). A and D utilize 2:1 DOPC:DOPG, B and E utilize 1:1 DOPC:DOPG, and C and F utilize 1:2 DOPC:DOPG. A-C lack salt or buffer in the aqueous while D-F include 10 mM HEPES, 0.1 M NaCl, 0.29 M sucrose (IA) or glucose (OA). We note that dewetting universally occurred, however no meaningful amount of GUVs survived the process intact.

Lipid compositions, salt, and flow speed influence dewetting dynamics in a context-dependent manner. In salt-free conditions, lipid concentration and flow speed act to impede dewetting dynamics, which contrasts with earlier reports showing that increasing lipid concentration and shear enhance dewetting^30^. While the dewetting results were classified into binary states, we observed quicker dewetting at low lipid concentrations and flow speed. The flow rate and lipid concentration dependency suggest that the Marangoni effect is at play. High concentrations of P188 and lipid could generate stronger Marangoni stresses, as fluid flow coupled to surfactant-laden droplets can induce surface tension gradients^54,58,72–73^. This may explain the failure to dewet at higher speeds with higher lipid concentrations (Figures 4A-C).

The lipid charge, which has not been explored, also showed a negative influence on dewetting in the case of no salt and buffer with 5% P188. In other words, conditions with a higher composition of negative lipids exhibited slower dewetting. Additionally, higher lipid charge showed a dynamic flow speed dependency: higher speeds impeded dewetting in conditions with the greater negative charge (1:2 DOPC/DOPG and 1:1 DOPC/DOPG), whereas the lowest negative charge (2:1 DOPC/DOPG) at the lowest lipid concentration dewetted on-chip at all flow speeds tested (Figure 4C). These results suggest that increased lipid charge enhances Marangoni effect in the absence of salt.

In contrast, the presence of salt and buffer (Figure 4D-F) counteracts the trends for lipid concentration and charge observed in no-salt conditions and reduces the impact of flow speed. Only the highest concentration of 35 mg/mL and the most negative charge (1:2 DOPC/DOPG) was able to dewet on-chip at all the flow speeds tested (Figure 4D). This trend reversal may be explained by the ability of salt to decrease lipid mobility^74,75^, increase the viscosity of the aqueous phase, and enhance membrane rigidity and thickness, owing to strong electrostatic interactions with lipid head groups, which dampen the Marangoni effect. These changes complicate the kinetics of dewetting that our results have revealed, making it difficult to predict by only considering the spreading coefficient. These findings underscore the greater complexity of on-chip dewetting dynamics than previously understood, going beyond the traditional thermodynamic equilibrium-based predictions.

### Dewetting Dynamics at Low OA Surfactant Concentration Without Glycerol

In contrast to the 5% P188 conditions, 0.1% P188 without glycerol produced intact GUVs but often failed to complete dewetting on-chip, with partially dewetted DE droplets persisting off chip for several days. Centrifugation post on-chip dewetting facilitated complete dewetting under certain conditions, broadening the operational window without requiring extensive optimization. We categorized the outcomes at 0.1% P188 into four categories (Figure 5): 1) on-chip full dewetting, 2) off-chip full dewetting, 3) partial dewetting on chip but dewetting completed after centrifugation, and 4) partial dewetting persisting even after centrifugation. The failure to dewet at certain conditions aligns with the trend implied by the surface tension measurements, which indicate that lower P188 concentrations in OA increase the surface tension γ_23_, thereby reducing the separation force.

**Figure 5.**
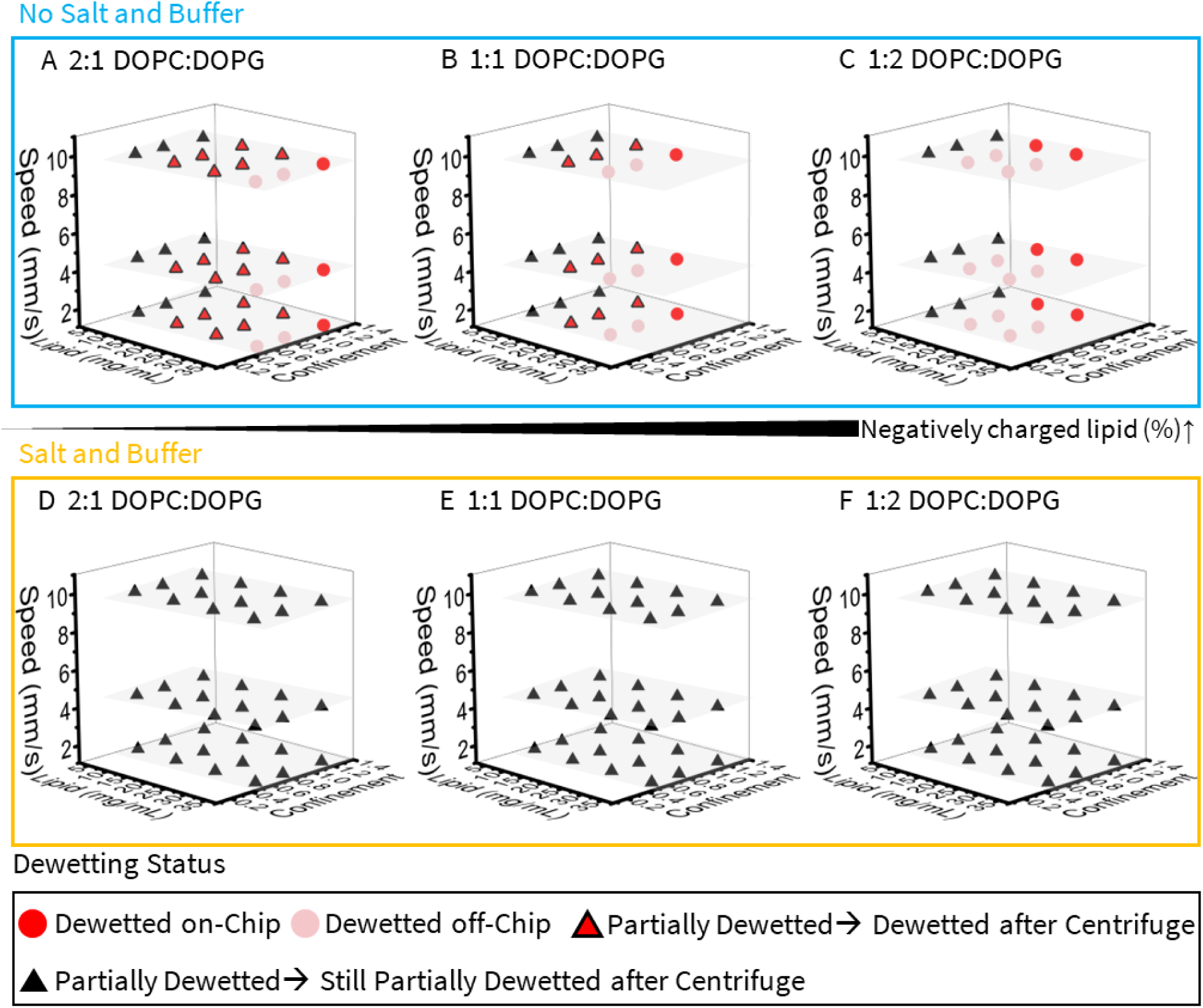
Dewetting results at low OA surfactant concentration (0.1% P188). Similar to Figure 4, A-C are for no salt or buffer in the aqueous while D-F contains 10 mM HEPES, 0.1 M NaCl, 0.29 M sucrose (IA) or glucose (OA).The four on-chip dewetting status are categorized as follows: 1) DE dewetted and produced intact GUVs within the dewetting-chip, 2) a fraction of DEs dewetted within the maximum on-chip time of ∼30 s but the remainder slowly dewetted off-chip, 3) DEs that partially dewetted on-chip or passively off-chip could be successful dewetted after centrifugation, 4) DEs did not dewet even at 17000 × *g* for 1 min.

Consistent with 5% P188, confinement greater than 1 was required for on-chip dewetting. However, we observed a significant difference in the dewetting dynamics for the 0.1% and 5% P188 conditions. The flow speed effect at high confinement diminished, and the dewetting time was substantially reduced. Specifically, on-chip dewetting with 0.1% P188, when observed, always occurred within the first 10 seconds (Figure 7), whereas dewetting with 5% P188 took considerably longer. Table S1 highlights a 10-fold decrease in dewetting time at 0.1% P188 compared to 5% under identical conditions. This result suggests that Marangoni effect was weaker due to significantly lower surfactant concentrations. Even in cases of partial dewetting, the transition from DE to the partially dewetted state occurred rapidly under higher confinement (Figure S4). These results demonstrate that droplet confinement (confinement >1) can enforce the dewetting process, as noted in previous studies.

In the absence of salt (Figure 5A-C), lipid concentration and negative charge synergistically enhance the dewetting success. At all charge compositions, the lowest lipid concentration, 5 mg/mL, failed to dewet even over several days off-chip or following centrifugation. At 25-35 mg/mL, complete on-chip dewetting was achieved while the intermediate concentrations required centrifugation post on-chip dewetting to achieve full dewetting. These results align with previous studies demonstrating that increasing lipid concentration was conducive to dewetting^30^. Higher lipid negative charge composition further promoted dewetting; for example, 35 mg/mL lipid was necessary for the lowest amount of charge (33% DOPG) while 15 mg/mL was sufficient at 67% DOPG. Overall, higher lipid concentrations and more charge are critical for successful on-chip dewetting under restricted P188 conditions.

Centrifugation drives separation of GUVs from partially dewetted droplets, expanding suitable conditions for GUV assembly without optimizing on-chip dewetting. Centrifugation resulted in four outcomes (Figure S9): 1) mostly intact GUVs, 2) a mix of intact and fragmented GUVs, 3) GUVs with droplets exhibiting incomplete dewetting, 4) most droplets remaining partially dewetted with fragmented vesicles. Figure 6 shows the size distribution and counts of fully dewetted GUVs after centrifugation, excluding partially dewetted vesicles. Starting with a similar number of DE droplets across all conditions, the GUV yield can be assessed based on their count. The size range of non-fragmented GUVs is characterized by using the on-chip dewetting results, where minimal GUV fragmentation is observed. Despite some GUV fragmentation, we obtained GUVs with a reasonable size distribution (CV<8%) using optimized conditions.

**Figure 6.**
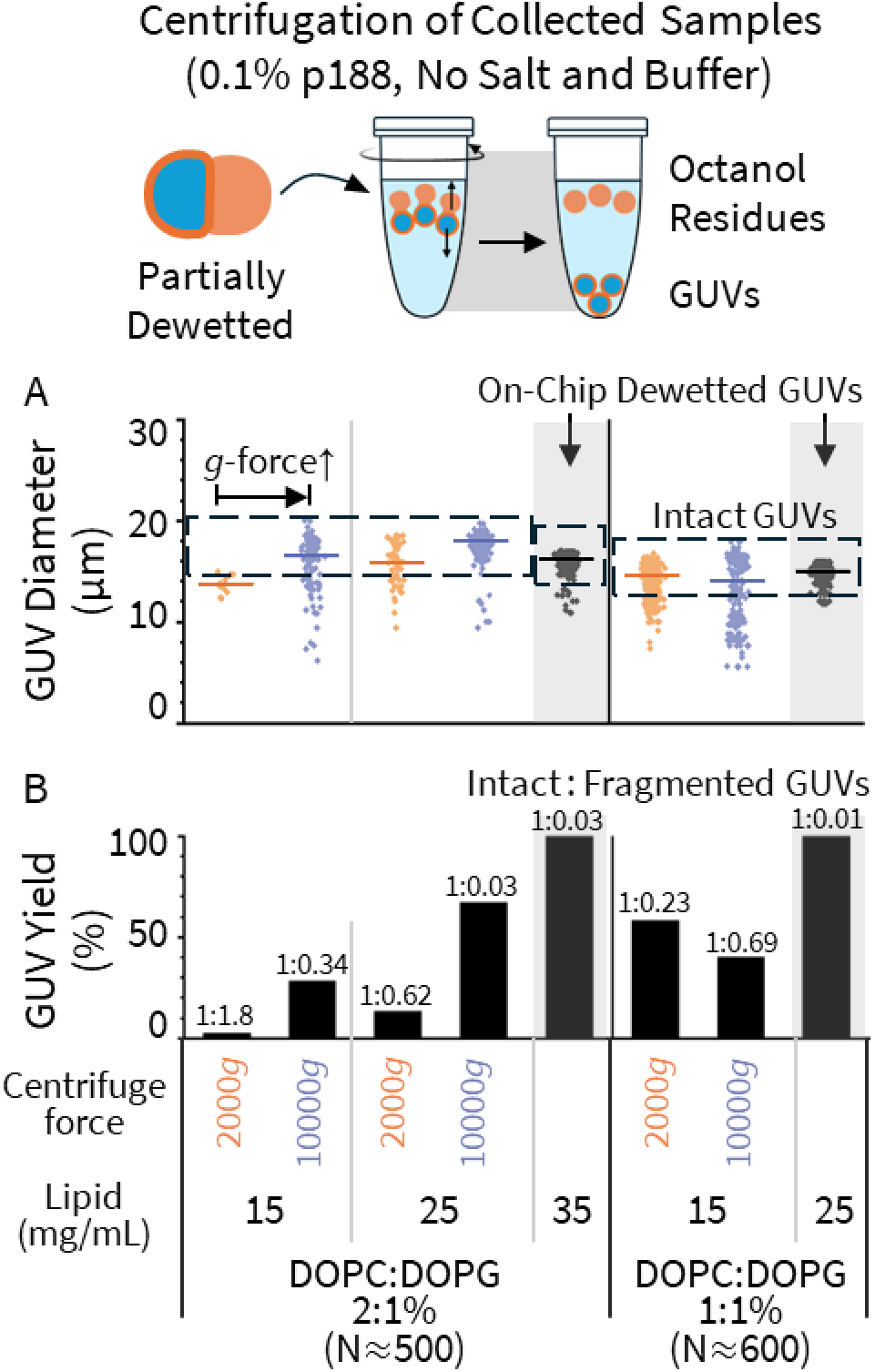
Centrifugation of partially dewetted GUVs with low P188 and no salt. All samples were centrifuged from a similar number of DE droplets. The bottom portion of the samples was imaged as GUVs sink during centrifugation. Fluorescently labeled GUVs were imaged and analyzed. (A) The diameter distribution of centrifuged or on-chip dewetted GUVs. The solid color-coated line across the population indicates the mean diameter of GUVs in the observed size distribution. The dashed black boxes represent the three-sigma cut-off to differentiate intact GUVs, which fall within the cut-off range, from fragmented GUVs, which exceed this range and are considered structurally compromised. These cut-offs are set at three-sigma of the size distribution of on-chip dewetted GUVs (1:1 DOPC:DOPG 25 mg/mL), where fragmentation was minimal. They are also adjusted to account for the relative size differences in the starting DE droplets, ensuring that the comparison remains consistent despite initial variations. (B) GUV yield (%) of centrifugation-driven dewetted GUVs relative to on-chip dewetted GUVs, where full dewetting with minimal GUV fragmentation was observed. The total number of intact and fragmented GUVs were counted and normalized to the corresponding counts in on-chip dewetted samples for specific lipid compositions (∼500 for 2:1 DOPC:DOPG and ∼600 for 1:1 DOPC:DOPG). As centrifugation began with a similar number of DE droplets, GUV counts were used to compare the yields of centrifugation and on-chip dewetting. It was not feasible to determine the exact number of vesicles each GUV fragmented into; therefore, both intact and fragmented GUVs were included in the yield calculation, with their ratios specified above the column. The specific centrifugation conditions, lipid combinations and concentrations are listed.

**Figure 7.**
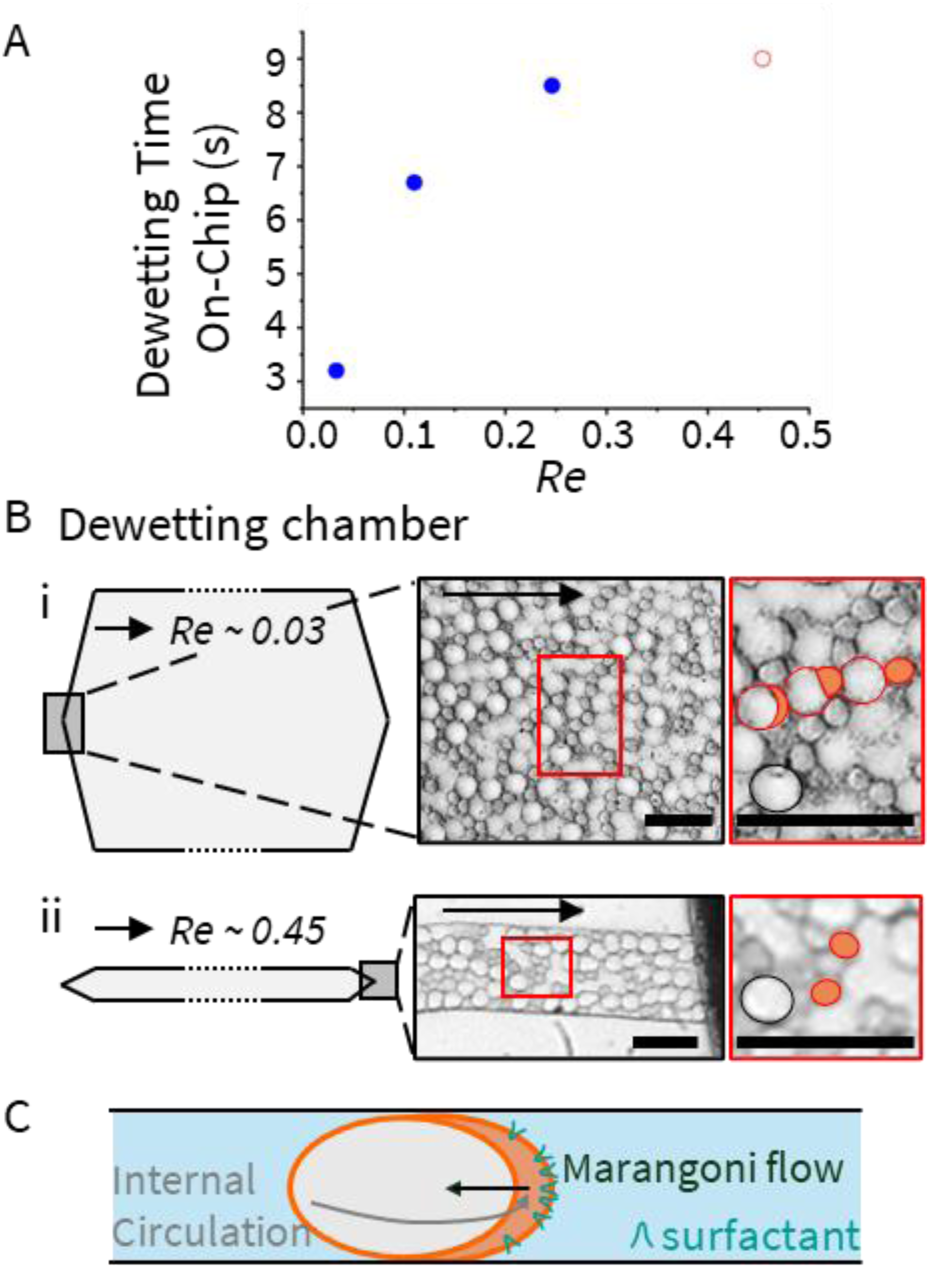
The effects of flow speed on dewetting for 2:1 DOPG:DOPC at 25 mg/mL, 1.2 confinement, 0.1% P188 in the OA, and no salt. A) Time required to achieve complete dewetting on chip, which is also tabulated in Table S2. Note that the red ring represents the maximum time on-chip at that speed as complete dewetting was not achieved like for the three lower speeds.

Effective completion of dewetting upon centrifugation depends on both lipid compositions and centrifugal force. High lipid concentrations and negative charge require a milder g-force for complete dewetting, aligning with the observed trends favoring on-chip dewetting (Figure 5). These findings suggest that centrifugation helps overcome kinetic barriers, with milder conditions effective for lower kinetic barriers. At 5 mg/mL of all lipid combinations, even the maximum centrifugal force (17000 × *g*) failed to fully dewet the droplets (Figure 5). In contrast, 15 mg/mL of DOPC/DOPG (2:1) required 10000*g* for efficient GUV formation (Figure S9). At 25 mg/mL, dewetting was effective even at 2000 × *g*. Moreover, 15 mg/mL of higher charge DOPC/DOPG 1:1 was efficiently separable at 2000*g*, generating more intact GUV counts than the same concentration of DOPC/DOPG (2:1) at higher g-force (Figure 6). While higher g-force generally facilitates complete dewetting, excessive forces can cause GUV fragmentation, as observed with 15 mg/mL of DOPC/DOPG (1:1). Thus, careful optimization of centrifugation parameters is essential for optimal separation while maintaining GUV integrity.

The dewetting results completely change upon the inclusion of physiological buffer (10 mM HEPES) and salt (100 mM NaCl), as shown in Figures 5D-F. Notably, none of the conditions achieve complete dewetting, even after extensive centrifugation (Figure S11). These findings are consistent with similar observations made by others, showing partial dewetting under physiological ionic concentrations, such as 180 mM potassium glutamate, even with 1% P188^46^. In addition to increasing interfacial tensions, salt can slow dewetting by increasing the viscosity of the aqueous phase and reducing lipid mobility^74,75^. These effects hinder intermolecular shuffling and the expulsion of the oil, both essential for successful dewetting.

Another study on liquid-liquid phase separation of polymeric composite particles supports this speculation, showing that kinetic barriers, such as increased viscosity of liquid phase from solvent evaporation, can prevent the system from reaching its predicted thermodynamic equilibrium^76^. While we will address the challenge of salt further in the last section, we have identified parameter adjustments to favor dewetting without relying on glycerol or high concentrations of P188.

### Influence of Fluid Dynamics on Dewetting

Therefore, the datapoint for the highest speed represents a lower bound on the on-chip dewetting time, which is >9 seconds. B) Morphology of double emulsions inside the dewetting at two different flow speeds. Arrows indicate flow direction. Red circles indicate droplets undergoing phase separation, orange circles highlight phase separated octanol droplets, and black circles represent non-dewetted DEs. Scale bars are 30 µm. i) 1.2 mm/s and lower shear. Most DEs dewetted shortly after entering the channel. ii) 19.2 mm/s and higher shear. Many double emulsions did not dewet at the outlet of the channel. C) Possible schematic representation of shear-induced surfactant redistribution and the Marangoni effect around DE droplet interface under confinement We examined the fluid dynamic effects on dewetting with 0.1% P188 and found that dewetting is more effective at lower flow speeds and thus lower shear rates, contrary to the previous assumption^26^. Figure 7A shows the correlation between on-chip dewetting time and the Reynolds number for the corresponding speeds under specified conditions. The three solid blue circles show that dewetting time increases with higher *Re*, all within the dewetting chamber. The red ring data point represents an on-chip dewetting time >9 seconds, at the highest speed of ∼19 mm/s or *Re* of ∼0.45, for which many DEs did not dewet at 9 seconds. Due to technical constraints, we could not test longer residence time at this speed, as attempts to increase distance led to high flow resistance and backflow. However, these results are sufficient to demonstrate the adverse effect of high speed on dewetting under confinement.

Droplet morphologies are notably different at different flow speeds (Figure 7B). At the highest speed (Figure 7Bii), DE and oil droplets show greater deformation. DE droplets also lack the characteristic octanol pocket observed at the front of partially dewetted state at lower speeds (Figure 7Bi). Despite the low surfactant (P188) concentration, surfactant accumulation at the front of droplets under strong forward internal circulation in a microfluidic channel may create a backward-facing surface tension gradient that opposes the octanol pocket formation in front (Figure 7C). This observation suggests that although high confinement accelerates the internal circulation and promotes transition toward the partially dewetted state^70^, it can further amplify the dynamic negative effects of high shear and/or stronger Marangoni stresses. These results highlight the influence of fluid dynamics on the dewetting process, particularly the inverse relationship between flow speed and dewetting time at high confinement.

In summary, 0.1% P188 altered the dewetting dynamics and achieved complete dewetting within seconds. In contrast, the original study with 5% P188 required several minutes, limiting flow rate and throughput to avoid impractically long channel lengths resulting in excessively high flow resistance. With 0.1% P188, shorter dewetting times and lower flow resistance, achieved by wider dewetting channels at slow speeds, work together to enhance throughput.

### Influence of adhesive energy of lipids to dewetting: Achieving intact GUV production Under Biorelevant Conditions with salt

At biologically tolerable P188 levels, salt and buffer hinder on-chip dewetting, but they are essential components of physiological systems, influencing protein structures, lipid-protein interactions, and membrane properties such as stability and packing. To address this, we examined the influence of lipid composition beyond lipid charge to achieve dewetting with salt and buffer. While on-chip dewetting remains possible, our focus was on systematically analyzing parameter effects rather than fully optimizing the on-chip process, thus we utilized the on chip dewetting followed by centrifugation to produce fully dewetted GUVs.

As shown in Figure 8, we tested different acyl chain saturation levels of both neutral and negatively charged phospholipids commonly found in biological membranes (Figure S12). A saturated lipid DPPC (16:0 PC) imparts higher rigidity and lipid packing compared to POPC (16:0,18:1 PC) and DOPC (18:1 PC) which contain one or two monounsaturated acyl chains, respectively. To mimic mammalian lipid compositions^77,78^, we chose 30% for POPG (16:0, 18:1 PG) and 20% for cholesterol.

**Figure 8.**
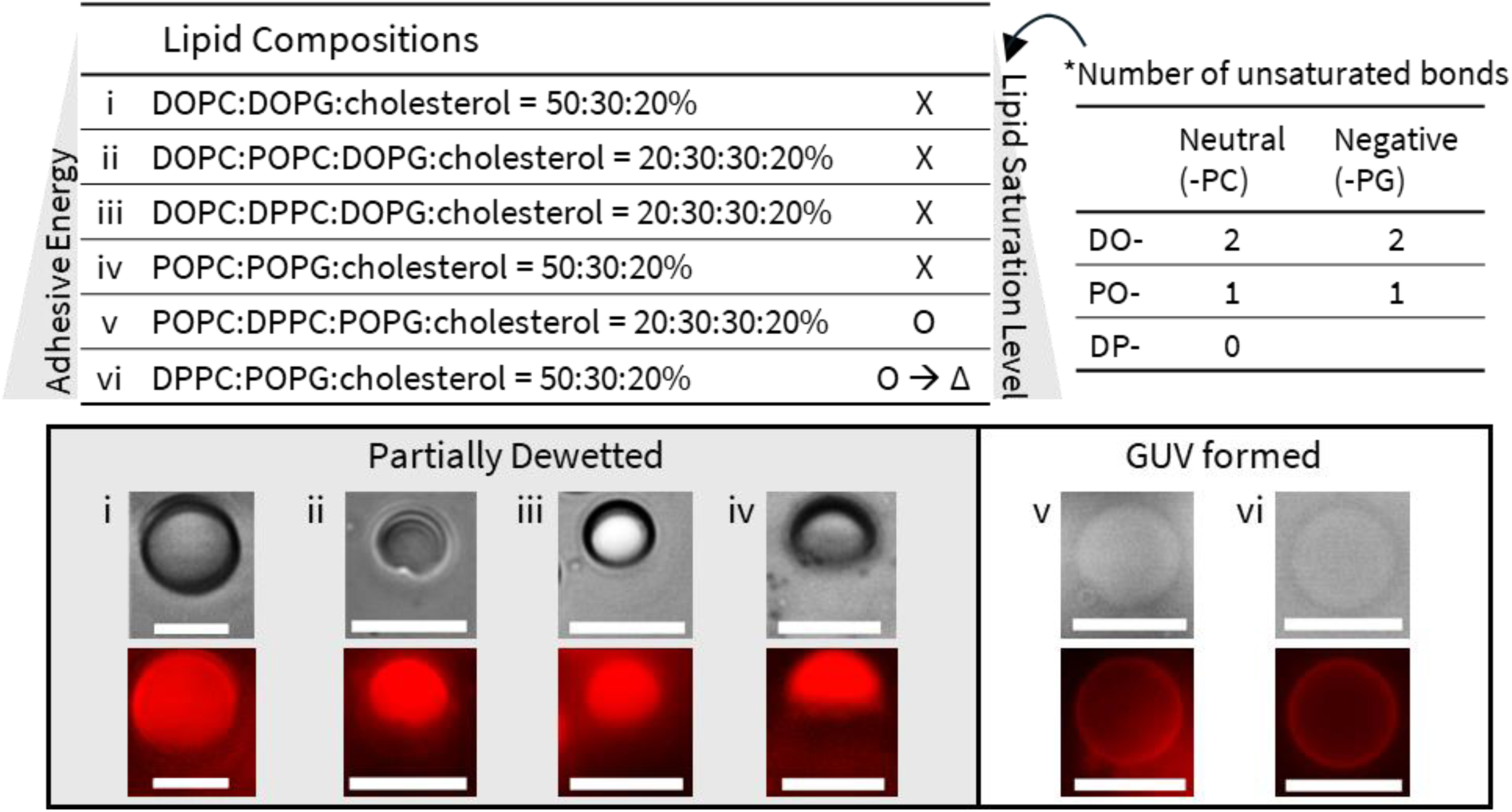
GUV generation under biologically relevant conditions. All conditions were prepared with 10 mM HEPES, 100 mM KCl, 290 mM glucose, 0.1% P188 in the OA with a lipid concentration of 25 mg/mL. The specific lipid compositions are detailed in the left table, while the number of unsaturated bonds in neutral and negatively charged phospholipids is indicated in the right table. The state of GUVs (2,000 × g, 1min) post-centrifuged are classified as follows. “X”: partially dewetted, “O”: completely dewetted, or “O→Δ”: completely dewetted but later fragmented. For each condition, both brightfield (upper panels) and fluorescent images (lower panels) of the post-centrifuged GUVs are shown. Scale bars: 10 μm.

We tested different conditions with lipids of increasing acyl chain saturation and found that higher saturation levels promote dewetting. For example, condition 1, the most disordered composition (DOPC/DOPG/cholesterol 5:3:2%), did not achieve complete dewetting. Replacing 60% of DOPC in condition 1 with POPC (condition 2) or DPPC (condition 3) did not enable complete dewetting. Even replacing all DOPC/DOPG in condition 1 with POPC/POPG (condition 4) failed. However, dewetting was successful by substituting 60% (condition 5) or 100% (condition 6) of POPC in condition 4 with DPPC. Unfortunately, condition 6 lacked long-term structural integrity, resulting in GUV fragmentation over days.

The effect of lipid acyl chain saturation can be understood through free energy minimization and the intermolecular forces at play. In droplet interface bilayer (DIB) formation, the oil phase is expelled as adhesion energy between the hydrophobic tails of the lipids increases, as described by the Young-Duprè equation^79,80^ (Figure S13):

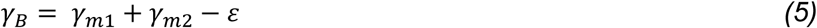

where γ_*B*_ is the bilayer surface tension, γ_*m*1_ and γ_*m*2_ are the surface tensions between the middle oil phase and each of the inner and outer aqueous phases, and ε is the adhesion energy that encompasses attractive and repulsive interactions. For DE dewetting, γ_*B*_ corresponds to γ_13_ (Figure 1), and the equation (5) adapts to *S*_2_ = - ε. Thus, bilayer formation is more favorable with higher adhesion energy, which is enhanced by tighter packing in fully saturated lipids. This perspective aligns with our observations: none of the tested conditions involving DOPC achieved full dewetting under biologically relevant conditions, likely due to its poor packing and low adhesion energy. In addition, disordered packing would retain more octanol in our DE droplet system, further hindering tighter lipid packing and increasing interfacial tension^81^.

Indeed, DOPC is notably challenging for DIB formation^82^, whereas DPhPC, a 16-carbon and fully saturated lipid originating from archaea, is commonly used. Although DPhPC differs from DPPC by methyl substitutions on its acyl chains, both pack more tightly than DOPC. Cholesterol at 20 mole% fraction, shown in DIB studies to maximize adhesion energy within 0-30% range^79^, supports its use in our experiments.

Conditions 5 and 6, which correspond to the highest saturation levels, were the only conditions to dewet, supporting the need to increase lipid hydrophobic interactions and increase adhesion energy. This result also supports our earlier observations that higher lipid concentrations promote dewetting, possible due to increased adhesive energy. A comparable mechanism was observed in the polymersome formation from double emulsions, where dewetting is driven by depletion interaction caused by excess polymer^83^. This excess polymer increases the adhesion energy, and faster dewetting is correlated with increasing polymer concentration in the solvent.

Increasing adhesion energy with higher fractions of saturated lipids, as in condition 6, may lead to fragmentation due to membrane asymmetries from microdomain formation (Figure S14). Fully saturated lipids such as DPPC^84^ or DSPC^85^ can phase separate into ordered domains, with a transition temperature of DPPC (41°C) contrasting with DOPC (−17°C) and POPC (−2°C). A previous study on ternary DPPC/DPPG/cholesterol compositions also show that the microdomain formation coupled with charged lipids and cholesterol can lead to pore formation and result in membrane morphology changes^86^. In condition 6, GUVs have non-spherical morphologies (Figure S14A) indicating asymmetric surface stresses^86^ that could lead to differential surface stresses and rupture. Another potential mechanism is GUV deflation due to pore formation, with fragmentation (Figure S14B) or division following from both the excess surface area and the phase separation^87^, though this needs further study. Compared to the condition 6, condition 5 contains lower DPPC and higher POPC molar fractions, which is likely to yield smaller domains and more stable GUVs^84^. Overall, these findings emphasize how lipid physical properties play a critical role in successful GUV dewetting as well as GUV stability, influencing factors such as adhesive energy and microdomain formation. These results serve as a guideline for optimizing lipid compositions to achieve desired experimental outcomes, specifically the production of stable GUVs under biologically relevant conditions.

## Conclusion

The bottom-up assembly of synthetic cells using GUVs offers an excellent opportunity for harnessing and studying both natural cellular processes and engineered ones, but a better understanding of the physical processes for generating these chassis is needed to fully realize the advantages promised by the double emulsion approach that is increasingly being relied upon. To this end, we conducted the first systematic investigation into the parameters influencing the dewetting dynamics of GUV production. By separating the droplet-making and dewetting modules, we could independently control confinement and flow dynamics and implement simple and robust fabrication methods. Our results showed how flow speed, droplet confinement, lipid type/composition/concentration, surfactant concentration, and added centrifugal forces should be tuned to achieve greater throughput and control. Specifically, a slow speed, high droplet confinement, elevated lipid concentrations/lipid charge, and the inclusion of saturated lipids promote the dewetting dynamics under biologically relevant conditions by influencing surface tensions and fluid dynamics. We show that it is possible to attain stable GUVs in biologically relevant conditions challenged by salts and buffers while eliminating the need for the widely used high glycerol and P188 concentrations, with the former interfering with reactions^31,43^ and the latter disturbing biological interactions and impairing cell functions. By integrating biocompatible conditions with the new chip design, our method achieves a throughput improvement by orders of magnitude. Given the multidisciplinary^5,14,22^ nature of GUVs for the study of biophysics, biochemistry, and biomedical applications, we hope our results will help to improve the robustness, reliability, and reach that droplet microfluidics-based GUV platforms can offer.

## Acknowledgements

This work was carried out in part in the Materials Research Laboratory Central Research Facilities, University of Illinois. The authors acknowledge the start-up funds from University of Illinois at Urbana-Champaign.

## Notes

### Competing Interest Statement

The authors have declared no competing interest.

